# Constrained Body Mass Evolution and Decoupled Morphological Rates in Plesiosaurs

**DOI:** 10.64898/2026.06.24.734298

**Authors:** Ruizhe Jackevan Zhao, Chi Zhang

## Abstract

Body size, through its links to various physiological traits, has often been hypothesized to influence evolutionary rates. Negative body size-rate correlations have been reported in the morphological or molecular evolution of several extant vertebrate groups, including mammals, birds, reptiles, and teleost fishes. In this study, we estimated body masses for 89 species of plesiosaurs, a clade of Mesozoic aquatic reptiles, and found that their body size evolution conforms to a three-regime Ornstein-Uhlenbeck process, indicative of constrained evolution. Rates of morphological evolution, inferred using the skyline fossilized birth-death process and the variable-rates model, show minimal support for a correlation with body size in this clade. Our results thus serve as a counterexample, suggesting that the negative body size-rate relationship is not a universal vertebrate pattern, but rather a trend restricted to certain lineages.

## RESULTS

### Phylogenetic analyses reveal heterogeneous evolutionary rates

We employed Bayesian tip dating under the skyline fossilized birth-death (SFBD) process^1,2^ to reconstruct the plesiosaur phylogeny. The 50% majority-rule consensus tree (Figure S1) recovers a topology largely consistent with previous studies^3,4^. Tip-dating results place the origin of Plesiosauria at approximately 221.22 million years ago (Ma), with a 95% highest posterior density (HPD) interval spanning from 230.74 to 214 Ma (late Carnian to Norian). A large polytomy exists among Rhomaleosauridae, Pliosauridae, Plesiosauroidea, and some Early Jurassic species, suggesting unresolved interrelationships among the early-diverging plesiosaurs. Polytomies also occurr within Rhomaleosauridae, Thalassophonea, Leptocleidia, and Elasmosauridae, supporting previous findings that the topology within clades remains poorly resolved.^5–10^

The unpartitioned character matrix yields elevated branch-specific rates at the origins of Thalassophonea, Cryptoclidia, and Polycotylidae, as well as among basal Plesiosauroidea (Figure 1A). The partitioned analysis (craniodental, axial, and appendicular) reveals a broadly consistent pattern, with elevated rates concentrated at the origins of certain major clades (Figures 1BCD). This pattern suggests that clades such as Thalassophonea, Cryptoclidia, and Leptocleidia underwent extensive and rapid morphological change during their early divergence, after which morphology remained relatively stable within each lineage.

The SFBD model accommodates piecewise variation in macroevolutionary parameters across predefined time bins.^1,12^ The comparatively low extinction rates observed during the Late Triassic (Figure 1F) facilitated a steady accumulation of lineages, culminating in high taxonomic diversity by the Early Jurassic.^13^ This pattern is well resolved in the fossil record owing to the high sampling rate characteristic of the Early Jurassic (Figure 1H). From the Middle Jurassic through the Turonian, plesiosaur evolution was governed by a dynamic of elevated speciation coupled with high extinction rate (Figures 1EF). After the Turonian, the net diversification rate dropped to its lowest level in plesiosaur evolutionary history (Figure 1G). This decline coincides with the extinction of pliosaurids^14^ and a marked reduction in the phylogenetic diversity of polycotylids.^15^ Elasmosauridae represents the sole plesiosaur clade that continued to flourish beyond the Turonian (Figure S1).^16,17^

### Three-regime Ornstein-Uhlenbeck mode of body mass evolution

We compiled a body mass dataset by combining volumetric-density estimates from Zhao^18^ and applying the body mass equations established therein. The dataset comprises 89 plesiosaur species, each represented by an osteologically mature individual (*sensu* Araújo and Smith^19^). Figure 2A shows body mass estimates mapped onto the branches of the pruned maximum a posteriori tree, revealing that large body mass evolved independently in thalassophoneans and elasmosaurids. Throughout their evolutionary history, most other plesiosaur clades maintained relatively modest body masses (Figure 2B).

We used a likelihood-based approach to compare the fit of evolutionary models to logtransformed body mass data (see STAR Methods). Among single-regime models, the Brownian motion (BM), early burst (EB), rate trend, and mean trend models yielded similar sample-size corrected Akaike Information Criterion (AICc) values, indicating indistinguishable performance (Figure 2C). The Ornstein-Uhlenbeck (OU) model provided the best fit within this class. We also fitted three-regime Brownian motion (BM3) and Ornstein-Uhlenbeck (OU3) models to the data, and the OU3 model achieved the best overall fit.

Because likelihood-based model comparison can erroneously favor the OU model when applied to small datasets,^20^ we partitioned the dataset into three subsets (Thalassophonea, Elasmosauridae, and the others) to further compare the BM and OU models using a Bayesian approach (see STAR Methods). The posterior probabilities and the calculated Bayes factors indicate that the OU model significantly outperforms the BM model when fitted to each subset (Figure 2D). We then summarized the posterior distributions of the OU parameters for each subset. The mean selective optimum (*θ*) for the “others” subset is around 407 kg, which is substantially smaller than those for Elasmosauridae and Thalassophonea (Figure 2E). Regarding the selection strength (*α*), the median value for Thalassophonea is lower than those of the other two subsets, and *α* reaches zero in many iterations (Figure 2F). Elasmosaurids, on the other hand, experienced the strongest selection on body mass among the three subsets.

### Neck length disparity in relation to body mass

We performed a cluster analysis based on relative neck length in plesiosaurs (Figure S2). The results indicate that neck length can be categorized into two groups (“long” and “short”), and bootstrapping supports the robustness of this dichotomy. We refrained from using the traditional terms “plesiosauromorph” and “pliosauromorph” here, because such empirical dichotomy^21^ differs somewhat from the clustering result.

**Figure 1:**
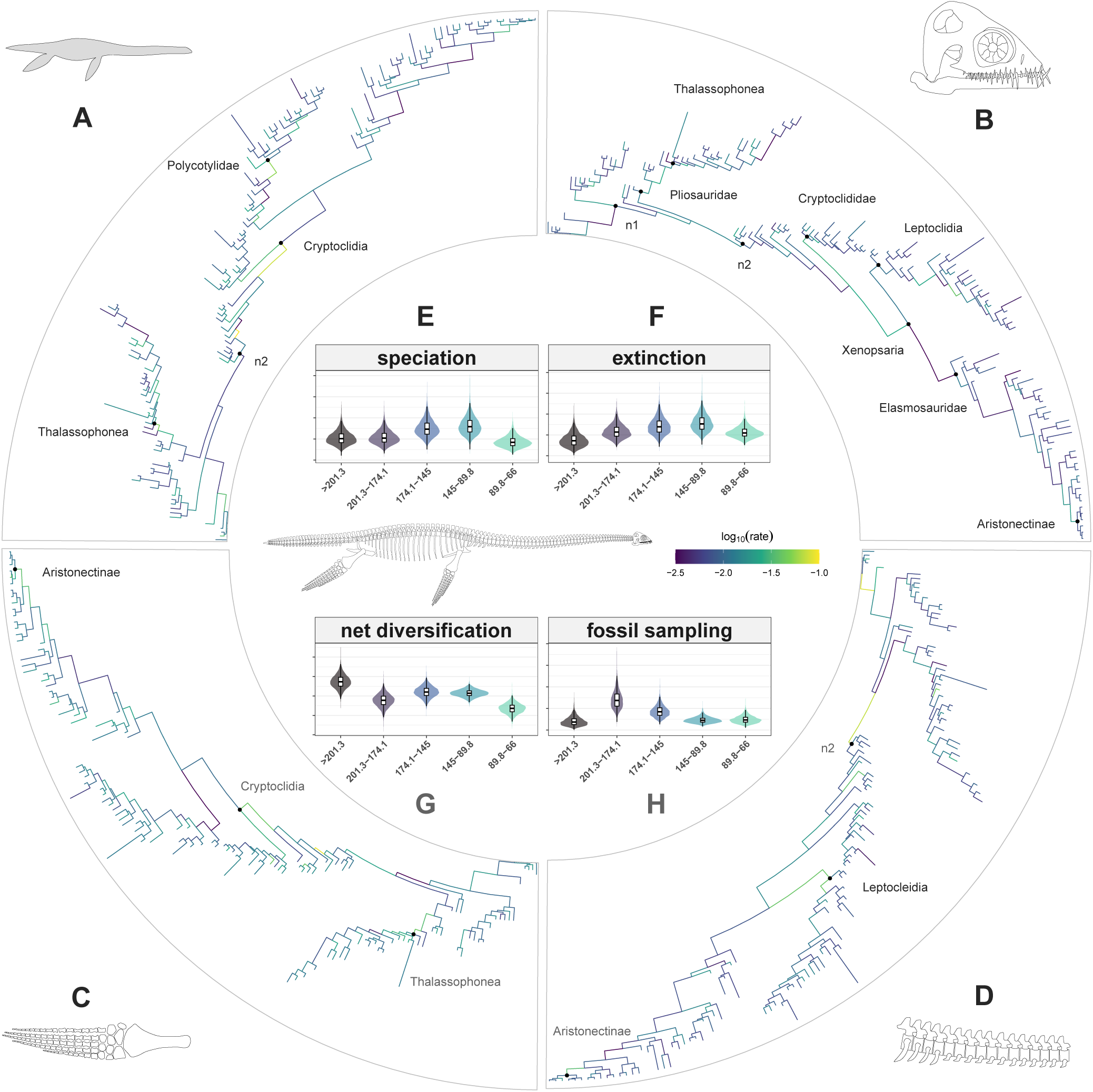
Results of Bayesian phylogenetic analyses based on the skyline fossilized birth-death model. (A-D) Rates of morphological evolution mapped onto the maximum a posteriori tree, computed from the (A) unpartitioned data matrix and the (B) craniodental, (C) appendicular, and (D) axial partitions, respectively. (E-H) Variation in (E) speciation rate, (F) extinction rate, (G) net diversification rate, and (H) fossil sampling rate across the five time bins. To avoid visual distortion caused by extreme rate values, the color scale for log-transformed rates was clipped to the range of -2.5 to -1. Values outside this interval were truncated to the corresponding endpoint. Node abbreviations: n1, Plesiosauria; n2, Plesiosauroidea. Definition of the time bins: bin 1, from the origin of plesiosaurs to the end of the Triassic (*>*201 million years ago [Ma]); bin 2, Early Jurassic (201.3-174.1 Ma); bin 3, Middle to Late Jurassic (174.1-145 Ma); bin 4, Cretaceous up to the Turonian (145-89.8 Ma); bin 5, post-Turonian Cretaceous (89.8-66 Ma). The skull of *Abyssosaurus nataliae* was redrawn according to Berezin11. Silhouette and other skeletal reconstructions were created by R.J.Z.

**Figure 2:**
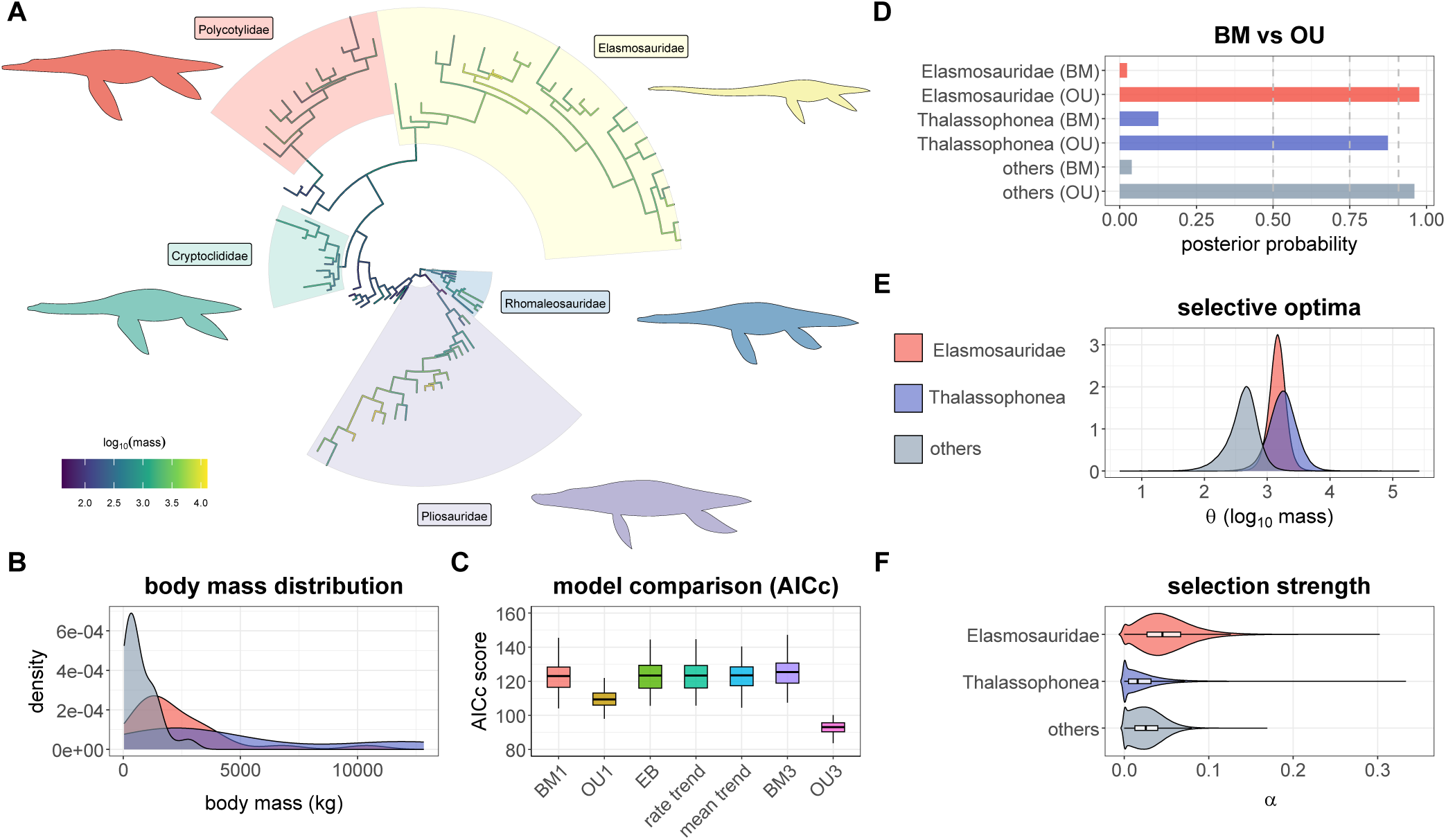
Body mass evolution of plesiosaurs. (A) Body mass estimates for 89 plesiosaur species mapped onto the branches of the maximum a posteriori tree. Lighter color indicates higher body mass. (B) Body mass distributions of the three subsets: Elasmosauridae, Thalassophonea, and the others. (C) Sample-size corrected Akaike Information Criterion (AICc) scores of the evolutionary models fitted to the log-transformed body mass data. Model abbreviations: BM1, single-regime Brownian motion; OU1, single-regime Ornstein-Uhlenbeck model; EB, early burst model; BM3, three-regime Brownian motion; OU3, three-regime Ornstein-Uhlenbeck model. (D) Posterior probabilities of the single-regime Brownian motion and Ornstein-Uhlenbeck models fitted to each of the three subsets. Dashed lines represent Bayes factors of 1, 3, and 10 (see STAR Methods). (E) Posterior distributions of the parameter *θ* of the OU models, representing the selective optima of the three subsets. (F) Posterior distributions of the parameter *α* of the OU models, representing the selection strengths of the three subsets. The legend in subfigure (E) also applies to (B) and (F). Plesiosaur silhouettes were created by R.J.Z, and are not to scale.

We performed a principal coordinate analysis (PCoA) on neck length, using the same distance matrix employed in the cluster analysis, and extracted the first principal coordinate (PCo1), which explained approximately 98.8% of the variance. We employed phylogenetic generalized least squares (PGLS) to test for an association between log-transformed body mass and neck PCo1 (Figure 3A). When the neck length dataset was analyzed in its entirety, none of the posterior trees examined rejected the null hypothesis, indicating no detectable correlation across all cases. For the long-necked subset, the null hypothesis was not rejected in 79% examined posterior trees, whereas for the short-necked subset, this proportion fell to 27%. These results indicate that the majority of trees support a correlation between log-transformed body mass and logtransformed neck length in short-necked taxa.

We investigated the relationship between neck disparity and body mass by dividing the neck dataset with the moving-window method (see STAR Methods). Bins are ordered by median body mass (Figure 3B). The error bars associated with the disparity estimates remained relatively uniform in length across all mass bins, indicating consistent estimation precision and statistical power throughout the body mass range. Elevated neck length disparity is evident in the second fixed bin and second overlapping bin (median body masses both around 640 kg). From the third fixed bin onward (median mass around 1255 kg), neck disparity shows a positive association with increasing body mass.

### Evolutionary rates decoupled from body mass

Our variable-rates model, fitted to neck PCo1 across 100 posterior trees (see STAR Methods), reveals a heterogeneous pattern of neck evolution (Figure 3C). The highest rates of neck length evolution were recovered within Cryptoclididae, where branch-specific rates were much higher than those estimated for other plesiosaurs. When Cryptoclididae is excluded from the dataset, additional peaks in evolutionary rate become apparent at the origin of Thalassophonea, along the stem of Leptocleidia, and at the origin of Aristonectinae. We used PGLS to evaluate whether rates of morphological evolution are correlated with body mass. Our analyses reveal that the vast majority of examined trees indicate no correlation between body mass and morphological rates for neck length (99%), the unpartitioned (78%), craniodental (84%), axial (89%), and appendicular (69%) datasets.

We reconstructed ancestral body masses at all internal nodes of the posterior trees under the OU3 model and calculated the mean mass for each branch (see STAR Methods). These branchwise body mass estimates were then used to evaluate the correlation between body mass and morphological evolutionary rate. The hexbin density plots (Figures 4A-D) reveal the posterior distributions of morphological rates across the body mass spectrum. For all character sets, branch rates display a unimodal, homoscedastic distribution. Most lineages concentrate within a dense, horizontal core, demonstrating a stable baseline rate of morphological evolution that is largely independent of body size.

We identified morphological rate bursts by extracting the top 5% of branch rates across each examined posterior tree, mapping their occurrence across the body mass spectrum (Figures 4E-H). Rather than a simple linear correlation, rate bursts exhibit complex, module-specific relationships with body size. The unpartitioned, axial, and appendicular datasets demonstrate multimodal burst distributions spanning distinct mass classes, with the unpartitioned and appendicular sets showing particularly diffuse density profiles. Conversely, craniodental bursts follow a unimodal distribution centered around 10^2.75^ kg. However, this peak is right-skewed, suggesting that rate bursts also happened in some lineages with large body masses.

**Figure 3:**
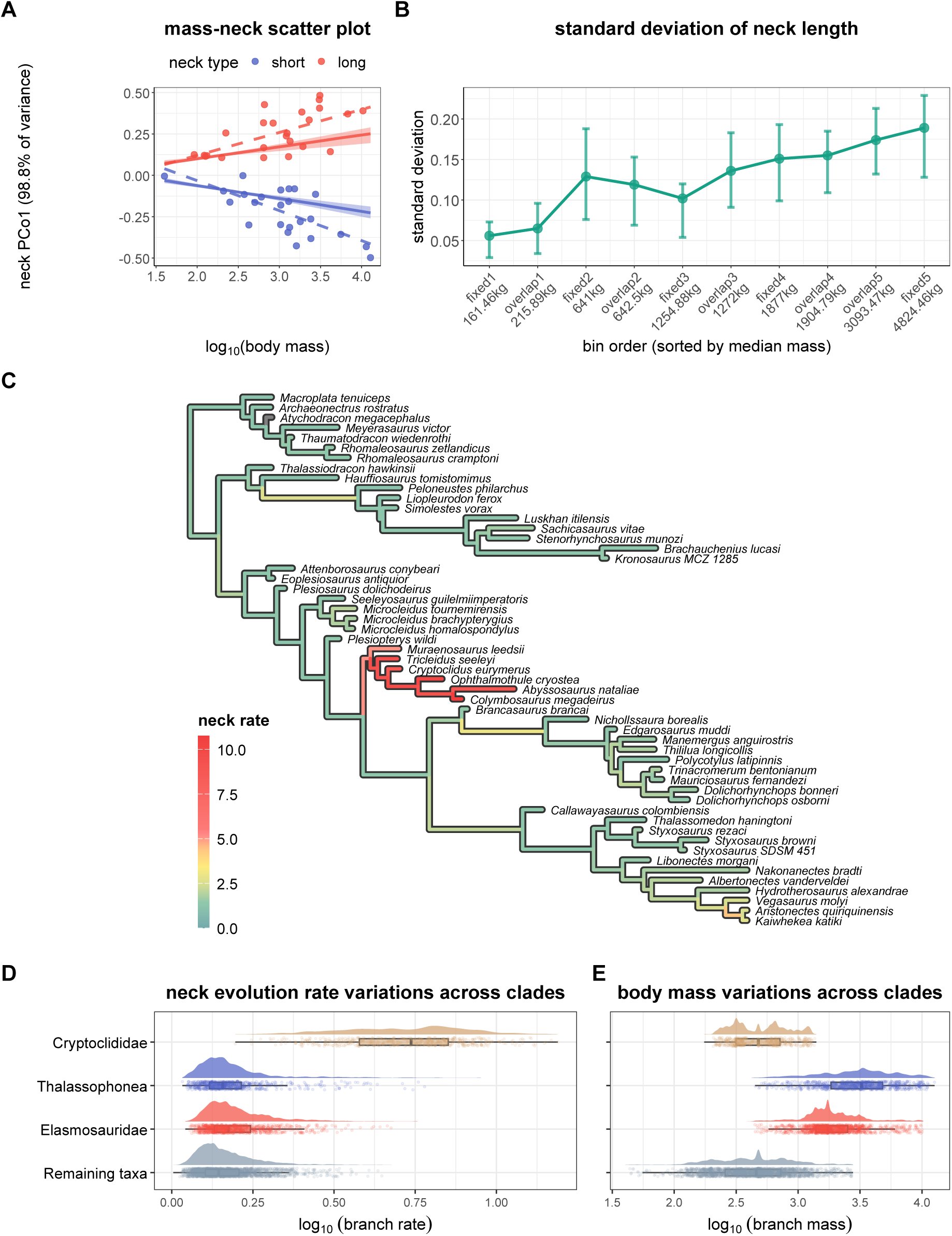
Plesiosaur neck length evolution in relation to body mass. (A) Scatter plot of log-transformed body mass versus the first principal coordinate (PCo1) of neck length, partitioned by the two clusters. Dashed line: ordinary least squares; solid line: mean coefficients of the phylogenetic generalized least squares (PGLS) models; shaded band: 95% PGLS confidence interval based on 100 posterior trees. (B) Pattern of variation in neck length across body mass bins. Error bars indicate the 95% confidence intervals based on the empirical bootstrap distributions of each bin (1,000 replicates). (C) Results of variable-rates model mapped onto one randomly selected posterior tree. (D) Raincloud plots showing the distribution of branch-specific rates of neck length evolution across four subsets. (E) Raincloud plots showing the distribution of branch mass across four subsets.

## DISCUSSION

Most published phylogenies of plesiosaurs were based on maximum parsimony methods.^3,9,17,22,23^ As demonstrated in many previous studies,^7,10,22–25^ plesiosaur phylogenies often exhibit large polytomies and are sensitive to parsimony weighting strategies, with alternative parameter settings producing markedly different topologies. Our 50% majority-rule consensus trees (Figure S1) indicate that the polytomy problem is not alleviated by the application of more complex Bayesian models. Notably, several clades that are relatively well resolved under maximum parsimony (e.g., Cryptoclididae^26,27^ and Polycotylidae^8,28^) also exhibit polytomies in the Bayesian framework. Our consensus tree also recovered a broad polytomy involving Rhomaleosauridae, Pliosauridae, Plesiosauroidea, and additional Early Jurassic species, reinforcing the idea that early plesiosaur relationships are still poorly understood.^3,23^ As evidenced by the discrepancies in PGLS results across posterior trees with varying topology and branch lengths, this study highlights the importance of accounting for phylogenetic uncertainty in comparative analyses.

Among the single-regime models, the OU process provided the best fit to plesiosaur body size evolution, whereas the remaining models tested were largely indistinguishable in their performance (Figure 2C). This suggests that body size evolution in plesiosaurs differs markedly from the early burst pattern seen in ichthyosaurs.^29,30^ The lack of support for an early burst also accords with the gradual morphospace occupation documented in Early Jurassic plesiosaurs and the smooth body size transition across the Triassic-Jurassic boundary.^13^ We also found that Cope’s rule^31^ can’t explain overall body size evolution in plesiosaurs, as evidenced by the low support for the mean trend model. This result is consistent with empirical observations, given that small-bodied plesiosaurs such as certain polycotylids persisted well into the Late Cretaceous.^15^ The reacquisition of an aquatic lifestyle imposes opposing constraints on body size in secondarily aquatic tetrapods: the need to limit heat loss sets a lower bound,^32,33^ while the challenge of meeting energetic requirements limits maximum size.^34,35^ This dual constraint is reflected by the body mass evolution of most modern aquatic mammals. Excluding a few clades such as the baleen whales, the macroevolutionary dynamics of aquatic mammals conform to an OU process, converging toward an adaptive optimum near 500 kg.^35^ Plesiosaurs resemble modern aquatic mammals in several physiological respects (e.g., viviparity^36^ and high metabolic rates^37^), and we note that the selective optimum recovered for small-bodied plesiosaurs (i.e., the “others” group in Figure 2) approaches the value estimated for extant aquatic mammals. It is therefore plausible that a similar trade-off mechanism also operated in plesiosaurs. In line with this, we recovered elevated neck length disparity within the second fixed bin and second overlapping bin (median body masses around 640 kg), suggesting that many plesiosaur clades may have explored distinct ecological niches while body size remained constrained to this general order of magnitude. Among the three partitioned groups, elasmosaurids experienced the strongest selection on large body mass (Figure 2F). A plausible explanation for this pattern lies in the energetic demands of underwater locomotion. Previous hydrodynamic work^38,39^ has demonstrated that the elongated neck of elasmosaurids generates considerable drag, and that a larger body size helps offset this cost by reducing the drag normalized by body mass. The body mass distribution of thalassophoneans is relatively flat (Figure 2B), and this group exhibits the weakest selection on body mass in the current partition scheme (Figure 2F). In contrast to elasmosaurids, the short neck characteristic of thalassophonean pliosaurs imposes fewer hydrodynamic constraints on body size evolution.^39^ Large body mass was attained by many species within both Thalassophonea and Elasmosauridae, and these two clades occupy opposite extremes of the neck length continuum.^21^ This pattern is effectively captured by our disparity analysis, which reveals a general increase in neck length disparity with increasing body mass in large species (Figure 3B). It is plausible that the independent acquisition of large body size in these two clades was permitted by fundamentally different feeding strategies. For instance, some thalassophoneans exhibit macrophagous ecomorphology,^40^ whereas elasmosaurids may have relied on their elongated necks for prey capture.^41^ However, such ecological interpretations remain speculative, as some fossilized stomach contents suggest that the potential prey of plesiosaurs may not correspond strictly to their feeding ecomorphology.^40,42,43^

**Figure 4:**
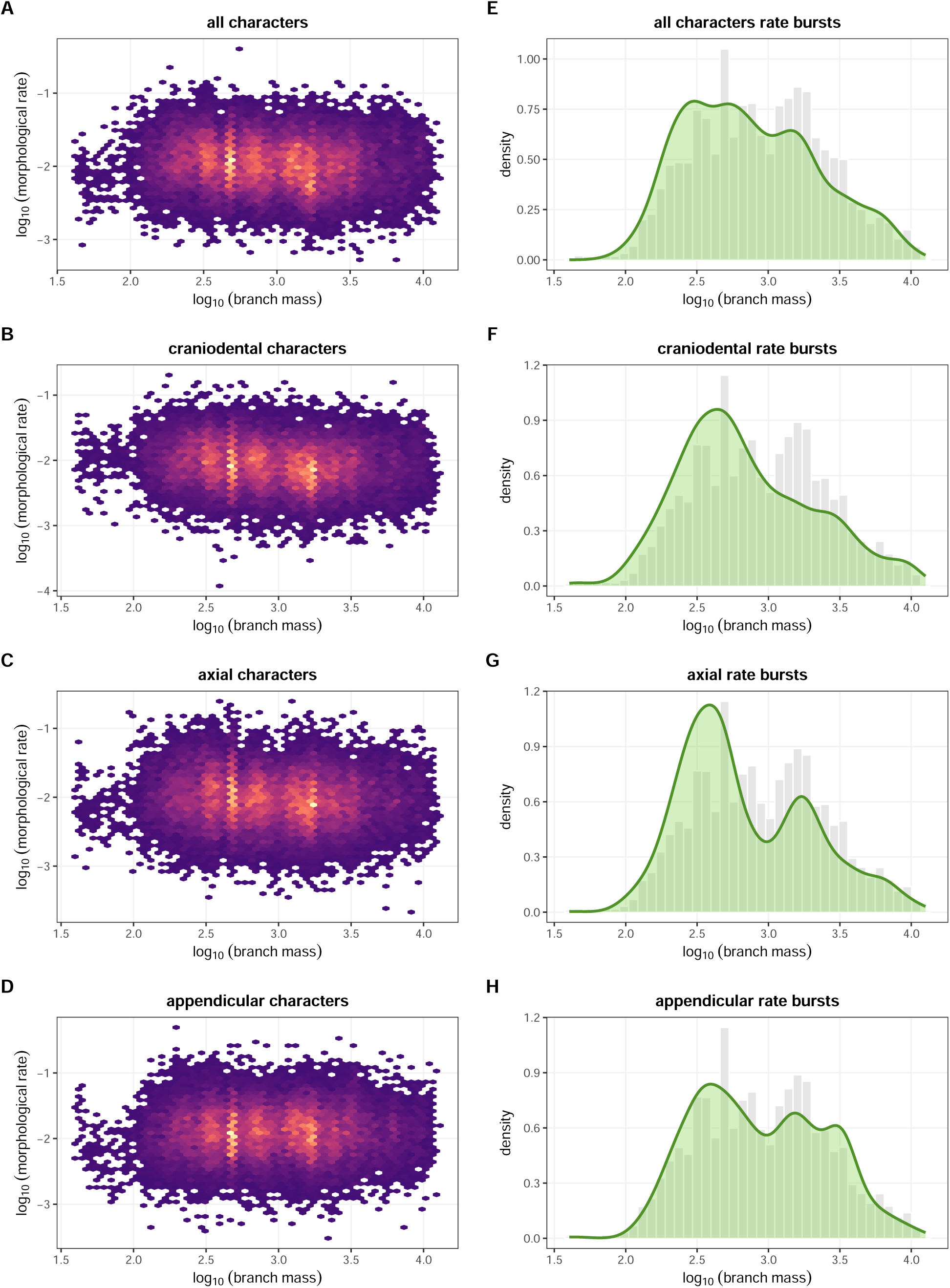
Plesiosaur morphological rates in relation to body mass. (A-D) Hexbin plots of the relationship between branch mass and morphological evolutionary rates for the (A) unpartitioned, (B) craniodental, (C) axial, and (D) appendicular datasets. Lighter color represents higher density in data distribution. (E-H) Body mass distributions for all phylogenetic branches (gray histograms) and rate bursts (green density plots) across (E) unpartitioned, (F) craniodental, (G) axial, and (H) appendicular datasets.

Our variable-rates model reveals that cryptoclidids exhibit extremely high rates of neck length evolution (Figure 3C). This finding is consistent with empirical observations, as cryptoclidids show remarkable variation in cervical count, ranging from approximately 27 vertebrae in *Tricleidus seeleyi* ^44^ to 60 in *Spitrasaurus wensaasi*.^45^ Beyond this clade, elevated rates of neck length evolution are typically restricted to the origin of new plesiosaur clades, such as Thalassophonea and Aristonectinae. A similar pattern, bursts of morphological innovation at the origin of new clades followed by relative stasis, was also recovered in our phylogenetic analyses (Figure 1), corresponding well to the evolutionary mode described as “quantum evolution”.^46^

Previous studies have widely assumed that animal evolutionary rates are influenced by various physiological factors.^47–51^ Because body size is often correlated with these underlying physiological metrics and is easier to measure,^47^ it has been utilized as a proxy in rate heterogeneity studies.^49,52^ A prevailing hypothesis is the “generation time effect”,^50,53^ proposing that smaller animals, due to their shorter generation times, can accumulate mutations more quickly per unit time, thereby responding more rapidly to natural selection. A negative correlation between body size and molecular evolutionary rates has indeed been documented in mammals,^49^ reptiles,^52^ and scombrid fishes.^54^ At the morphological level, birds^55^ and teleost fishes^56^ also exhibit a similar inverse relationship between body size and evolutionary rates. However, this negative correlation is not a strict biological law; rather, it may be an exception limited to certain extant vertebrate clades. For instance, molecular evolution in invertebrates displays heterogeneous patterns that are decoupled from body size.^57^ The impact of body size on vertebrate morphological evolution also takes on more complex forms. A relatively small body size subjects a terrestrial or flying animal to weaker mechanical constraints, which alters the mapping between form and function^58,59^, increases evolutionary lability,^60^ or permits the emergence of morphological innovations (e.g., the origin of avian flight^31,61^). Furthermore, when gravitational constraints are released through reacquisition of an aquatic lifestyle, the morphological evolution of marine mammals is profoundly altered, manifesting in the repatterning of evolutionary integration^62,63^ and the relaxation of selective pressures on certain anatomical structures.^64^

Across the datasets tested in this study, we generally found little support for a correlation between body mass and morphological evolutionary rate in plesiosaurs, and bursts in morphological evolutionary rates are not concentrated within specific body mass intervals, suggesting that rapid morphological shifts in plesiosaurs are not restricted to lineages of particular sizes. We propose that the buoyant aquatic environment possibly buffered plesiosaurs against the biomechanical penalties typically associated with large body size in terrestrial or flying vertebrates, thereby releasing their morphological potential. Furthermore, the K-selected life-history strategy of plesiosaurs^36^ might diminish the generation time effect. If extensive parental investment was a shared trait across the clade, the expected correlation between body mass and generation time might be disrupted. Although the exact reasons remain elusive, our results serve as a counterexample demonstrating that the negative correlation between body size and evolutionary rates does not apply to all vertebrates.

This study estimated plesiosaur morphological evolutionary rates based on neck length data and discrete character matrices; however, the influence of body size on plesiosaur evolution might not be reflected solely by shifts in morphological rates. As a natural extension of this work, shifting the perspective from evolutionary rates to plesiosaur ecomorphology and their internal patterns of evolutionary integration warrants future exploration. Tracing how shifts in body size rewire the evolutionary plasticity and covariation among skeletal structures will ultimately shed light on the broader pathways of morphological innovation following the gravitational release of a pelagic lifestyle.

## RESOURCE AVAILABILITY

### Lead contact

Requests for further information and resources should be directed to and will be fulfilled by the lead contact, Ruizhe Jackevan Zhao.

### Materials availability

This study did not generate new fossil materials.

### Data and code availability

- The data and code required to reproduce the analyses presented in this study is available at Zenodo (https://doi.org/10.5281/zenodo.20834674), which is a released version of the Github repository https://github.com/Pliosaurus-kevani/plesiosaur-size-evolution

## ACKNOWLEDGMENTS

This study is based on the undergraduate thesis of the first author, who thanks Andrew Orkney for his continuous guidance and discussions on phylogenetic comparative methods over the past three years. This work was funded by National Key Research and Development Program of China via grant 2023YFF0804502 to C.Z.

## AUTHOR CONTRIBUTIONS

Conceptualization, R.J.Z., and C.Z.; methodology, R.J.Z., and C.Z.; investigation, R.J.Z., and C.Z.; writing-–original draft, R.J.Z.; writing-–review & editing, R.J.Z. and C.Z.; funding acquisition, C.Z.; resources, R.J.Z. and C.Z.; supervision, C.Z.

## DECLARATION OF GENERATIVE AI AND AI-ASSISTED TECH-NOLOGIES

During the writing stage of this work, the authors used DeepSeek V4 in order to refine the wording and grammar of the manuscript. After using this tool, the authors reviewed and edited the content as needed and take full responsibility for the content of the publication.

## STAR METHODS

### Method details

#### Phylogenetic data

We employed Bayesian tip dating under the SFBD model^1,2^ to reconstruct the plesiosaur phylogeny. For extinct clades, this method requires a character matrix and stratigraphic information to perform phylogenetic analysis.^12,65^ We selected a published character matrix,^66^ which was extensively modified from Bensen and Druckenmiller^3^ by subsequent studies.^17,67–71^ It contains 290 characters and 130 operational taxonomic units (OTUs), which is one of the largest plesiosaur matrix to date. Most stratigraphic information for the OTUs was obtained from Madzia and Cau,^4^ with the range for *Ophthalmothule cryostea* modified following Roberts et al.^27^ Additional temporal data were retrieved from the Paleobiology Database (https://paleobiodb.org/).

The character matrix was divided into ordered and unordered categories. We applied the widely used^22,24,25,72,73^ scheme of 67 ordered characters proposed in Madzia et al,^74^ and further constrained character 155 to be ordered, following previous publications.^70,71^ We also conducted a partitioned analysis in which evolutionary rates were permitted to vary independently across three anatomical regions: craniodental, axial, and appendicular modules. Processing of the morphological characters were carried out in Mesquite.^75^

#### Body mass estimation

A recent study^18^ established a hybrid approach for estimating body mass in plesiosaurs. It obtained body mass estimates for multiple species through volumetric modeling and the cross-sectional method,^76^ then used them to identify skeletal proxies that effectively predict body mass. Among the metrics tested, dorsal centrum volume (mean centrum length *×* mean width *×* mean height) and trunk length were found to be the best predictors. Here we used ordinary least squares (OLS) equations based on these two variables to estimate body mass of plesiosaurs:

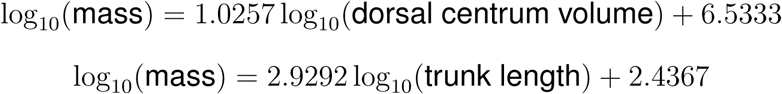

To expand the body mass dataset, we added species lacking dorsal vertebral data or measurable trunk length. Their body masses were computed using the full set of limb- and girdle-based OLS regression equations provided by Zhao^18^ (see table 2 in that study for all the equations), with the mean across all applicable formulae taken as the point estimate. For *Thaumatodracon wiedenrothi* and *Edgarosaurus muddi*, body mass was estimated using the OLS equation based on skull length and cervical count:

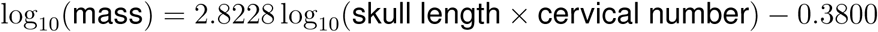

Additionally, some thalassophonean pliosaurs in the phylogeny are known mainly from skulls, with very limited postcranial material (e.g., *Pliosaurus westburyensis* ^77^ and *Pliosaurus almanzaensis* ^78^). To increase sample size, we estimated their body mass by scaling the skull-mass proportion of close relatives. We did not include juvenile specimens in our body mass dataset, as they may confound ecological constraints on body size in downstream analyses. We used the dataset from Araújo and Smith^19^ as a reference and included only individuals labeled as “osteologically mature” under the two-group (mature-immature) scheme. In total, our final body mass dataset comprises 89 plesiosaur species, surpassing those used in previous studies on plesiosaur body size evolution.^13,39^

### Quantification and statistical analysis

#### Phylogenetic analysis

The SFBD analysis enables the joint estimation of tree topology, branch lengths, and macroevolutionary parameters including speciation rates, extinction rates, and fossil sampling rates.^65,79^ This method allows these macroevolutionary parameters to vary piecewise across user-defined time bins.^1,2^ The evolutionary history of plesiosaurs was shaped by several extinction events: the Triassic - Jurassic,^13^ Early - Middle Jurassic,^80^ Jurassic - Cretaceous,^3,81^ Cenomanian - Turonian,^14,15^ and Cretaceous - Paleogene (K-Pg).^82^ Accordingly, we partitioned the plesiosaur evolutionary history into five time bins based on the extinction events. Since the occurrence times of the OTUs are uncertain, we allowed each tip date to vary randomly between its first appearance date (FAD) and last appearance date (LAD), following a uniform distribution. The matrix also includes several Triassic sauropterygians;^66^ accordingly, we incorporated these taxa when calibrating the root of the phylogeny. The root age was assigned a uniform prior distribution spanning from 247.2 Ma to 251 Ma. The former corresponds to the FAD of the oldest OTUs in the matrix (*Pistosaurus* skull and postcranium, and *Augustasaurus hagdorni* ).^4^ The latter is close to the beginning of the Early Triassic,^83^ substantially predating all OTUs included in the matrix.^4^ Because the current implementation of the SFBD model lacks an explicit parameter for the extinction of plesiosaurs, the algorithm would otherwise assume that unobserved lineages persist to the present day unsampled. To circumvent this issue, we shifted all occurrence ages forward by 66 million years, thereby treating the last plesiosaurs as effectively “extant” within the analysis. We fixed the extant sampling rate parameter to 1, assuming that the fossil record adequately captures plesiosaur evolutionary dynamics before the K-Pg extinction (e.g., the flourish of elasmosaurids^17^ and the decline of polycotylids^15^).

In addition to branch lengths, the tree topology also varies across iterations. Following Benson and Druckenmiller,^3^ we set *Yunguisaurus liae* as the outgroup. To reduce model complexity, we imposed topological constraints based on well-established monophyletic clades from previous studies: Rhomaleosauridae (excluding *Macroplata tenuiceps* ^84^ and *Anningasaura lymense* ^85^),^3,6,86^ Thalassophonea,^3,7,10^ *Seeleyosaurus guilelmiimperatoris* + *Microcleidus* spp.,^22,25^ Cryptoclididae,^26,27,87^ Leptocleidia,^28,88^ Polycotylidae,^8,28^ and Elasmosauridae.^9,17,71^ The phylogenetic positions of some early-diverging plesiosaurs, such as *Thalassiodracon hawkinsii*, *Attenborosaurus conybeari*, and *Stratesaurus taylori*, remain contentious and have varied in previous parsimony analyses under different weighting schemes.^22–24^ Accordingly, we did not impose topological constraints on these Early Jurassic taxa (as well as Pliosauridae and Plesiosauroidea), instead allowing their positions to vary across trees.

Character evolution was modeled under the Markov k-states variable (Mkv) model,^89^ which posits equal transition rates among states of a character. To capture among-character rate heterogeneity, we used discretized Gamma distribution with four categories following previous work.^90^ For the branch rates, we relaxed the constant rate assumption and employed a lognormal clock. Under this setting, evolutionary rates vary among branches and are independently sampled from a lognormal distribution. For the partitioned analysis, we unlinked the relaxed clocks assigned to the three skeletal modules, thereby decoupling their respective evolutionary rates.

The phylogenetic analysis was performed in RevBayes^91^ 1.3.2. We ran 500,000 generations of Metropolis - coupled Markov chain Monte Carlo (MCMCMC) iterations using four chains (1 cold and 3 heated chains) and four independent runs. To increase mixing efficiency among the chains, we employed a dual-swapping strategy. Swaps between adjacent chains were proposed every 10 iterations, and random swaps between randomly selected chains were proposed every 100 iterations. Before the focal analysis, we specified an initial 10,000 generations as burn-in and used the tuneHeat argument of the MCMCMC function during this stage to automatically adjust the temperature differences among the parallel heated chains. Trees and output files were sampled every 500 generations. For the partitioned analysis, we increased the number of generations to 1,000,000, as this substantially more complex model incorporates several hundred additional parameters.

We discarded the first 10% of samples as burn-in, and checked convergence of the posterior samples (consistent parameter estimates across runs and effective sample size *>* 200) using Tracer^92^ 1.7.2. All runs (both unpartitioned and partitioned) reached convergence. We subsequently summarized the maximum a posteriori tree and the 50% majority-rule consensus tree from the posterior sample of the combined runs for each analysis. To account for topological uncertainty in downstream analyses, we also randomly selected 100 posterior trees from each analysis.

#### Mode of body mass evolution

Given that large body mass evolved independently in thalassophoneans and elasmosaurids,^18^ we compared the fit of BM3 and OU3 models to log-transformed mass dataset against various single-regime alternatives, including BM, OU, early burst, rate trend (evolutionary rate changing linearly towards larger or smaller values), and mean trend (body mass having a trend towards larger or smaller values) models. To perform multi-regime model fitting, phylogenies with states mapped onto the branches are required. Because Thalassophonea and Elasmosauridae are both monophyletic clades^3,17^ in our partition scheme, we constructed a custom Markov model with three character states: Thalassophonea, Elasmosauridae, and all other plesiosaurs. In this model, transitions were permitted only from the “others” state to each of the two clades, thereby simulating their independent origins. All other possible transitions were forbidden. We performed stochastic character mapping using the make.simmap() function from the R package phytools^93^ version 2.5-2. We fitted the single-regime or three-regime BM and OU models using the mvBM() and mvOU() functions from the package mvMORPH^94^ version 1.2.1. We fitted the other single-regime models using the fitContinuous() function in the R package geiger^95^ version 2.0.11. Model fitting was replicated across the 100 randomly selected posterior trees, and the fit of all likelihood-based models was compared using the AICc values.

To further evaluate whether the OU model significantly outperforms the BM model in explaining body mass evolution within each subset, we split the dataset by regime and fitted the two single-regime models separately to each subset. It has been demonstrated that likelihood-based methods may mistakenly favor the OU model over the BM model, especially when the sample size is small.^20^ For robust inference, we employed a Bayesian method here. The OU model has the following form:

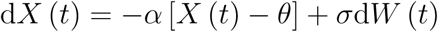

where the parameter *α* represents the strength of selection toward the optimum *θ*.^96^ When *α* equals 0, the OU model reduces to a BM model. Here we used a reversible-jump Markov chain Monte Carlo (rjMCMC) approach to compare the BM and OU models, which allows *α* to reach zero. After model fitting, we computed the Bayes factor (BF) of the OU model as:

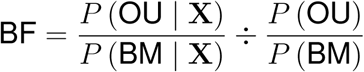

where X is the data set. We set equal prior probabilities for the BM and OU models, so the BF value is determined solely by the posterior probabilities. We implemented the model comparison in RevBayes^91^ 1.3.2. For each of the 100 trees, we ran 50,000 iterations, with the first 1,000 discarded as burn-in. We assessed convergence (effective sample size *>* 200) using the R package coda^97^ version 0.19-4.1, and the posterior distributions of all parameters were then summarized for visualization.

#### Neck length and body mass correlation

We used PGLS to investigate whether neck length in plesiosaurs is correlated with body mass. Large body size evolved independently in thalassophoneans and elasmosaurids, which had distinctly different neck lengths.^18,39^ Because PGLS assumes a linear relationship,^98^ we first performed a cluster analysis to determine whether plesiosaur neck lengths could be categorized into distinct types. We expanded the neck length dataset from Zhao^18^ by adding the ratio of neck length to trunk length as a new variable. Prior to the analysis, we log-transformed the continuous variables and then applied a z-transformation (so that each log-transformed variable had a mean of 0 and variance of 1). The dataset also contains a discrete variable (number of cervical vertebrae), so we computed the Gower distance matrix using the daisy() function from package cluster^99^ version 2.1.8.2 for the cluster analysis. We used the Ward.D2 criterion with 10,000 bootstrap replicates, implemented using the clusterboot() function from the R package fpc^100^ version 2.2-14. To evaluate the robustness of each cluster, this method generates a large number of bootstrap datasets and calculates the Jaccard similarity coefficient to identify the best-matching groups to the original clusters. A mean Jaccard coefficient greater than 0.85 indicates a highly stable cluster.^101^

We performed a PCoA based on the same Gower distance matrix, using the cmdscale() function from the stats package in R^102^ version 4.5.2. We then extracted the PCo1, which explains 98.8% of the variance. The dataset was divided into two neck types (long and short) based on the clustering result, and we performed PGLS between PCo1 and log-transformed body mass for each subset. We implemented the PGLS models using the phylolm() function in the R package phylolm^103^ version 2.6.5, with tree branch lengths transformed via Pagel’s *λ*.^104^ The PGLS models were replicated on the 100 randomly selected posterior trees.

We also partitioned the plesiosaur dataset into multiple bins by body mass and calculated the standard deviation of neck PCo1 within each subset. Since 73% of the examined trees indicated a correlation between neck PCo1 and body mass in short-necked taxa, we used the mean residuals from the 100 PGLS models as the proxy for neck length in this group. For long-necked taxa, we also took the mean residual, although 79% of trees indicated no correlation between neck length and body mass. This ensured consistency with the short-necked taxa: the expectation of the regression residuals is zero, while raw PCo1 values lack this property. To increase the resolution of our analysis, we applied a modified version of the moving-window method from Rothier et al.^59^ First, we partitioned the dataset into 5 fixed bins of increasing body mass, each with approximately the same number of samples. Then, we constructed overlapping bins by extracting samples from the adjacent fixed bins. The first overlapping bin draws all samples from the first fixed bin and half of the samples from the second fixed bin. Similarly, the last overlapping bin comprises all samples from the last fixed bin and half from the preceding bin. Each of the remaining overlapping bins takes all samples from the fixed bin with the corresponding index, plus one quarter of the samples from each of the two adjacent fixed bins. For each bin, we performed 1,000 bootstrap replicates and obtained the 95% confidence interval from the resulting empirical distribution (shown as error bars in Figure 3B).

#### Evolutionary rates and body mass correlation

We evaluated whether the rate of neck length evolution is correlated with body mass in plesiosaurs. To estimate branch-specific rates of neck evolution, we used neck length PCo1 obtained above as the variable, and employed the variable-rates model^105^ implemented in BayesTraits 4.0.0. By comparing the observed trait variation along each branch against expectations from a homogeneous Brownian motion, the model estimates branch-specific scalars that modify the background evolutionary rate.^105^ It employs a rjMCMC algorithm to automatically locate these rate shifts across the branches without prior assumptions. For each of the 100 randomly selected trees, we ran 12,000,000 iterations, discarding the first 2,000,000 as burn-in. Results were sampled every 10,000 iterations, and we summarized the mean branch-specific neck evolutionary rates in R. Convergence (effective sample size *>* 200) was assessed using the coda package^97^ version 0.19-4.1. To visualize the pattern of rate heterogeneity in the neck length evolution of plesiosaurs, we randomly selected a tree and mapped the evolutionary rates onto its branches using the ggtree package^106^ version 4.0.4.

To investigate whether the rate of neck length evolution is correlated with body mass, we extracted the morphological rates from the terminal branches and performed PGLS regression between log-transformed rates and log-transformed body mass. This analysis was replicated across all 100 posterior trees using the phylolm() function, with each tree rescaled according to Pagel’s *λ*. We observed that cryptoclidids exhibit extremely high rates of neck length evolution, and we explored whether this pattern is associated with distinctive body masses. Given that phylogenetic regression models can not account for autocorrelation among internal branches, we adopted a visualization-based approach to explore whether the distribution of neck evolutionary rates corresponds with that of body mass. We extracted the neck length evolutionary rates for all branches (both internal and terminal) and generated raincloud plots for Cryptoclididae, Thalassophonea, Elasmosauridae, and the remaining taxa. To estimate the mean body mass along each branch, we reconstructed ancestral states under the OU3 model using the estim() function in the R package mvMORPH^94^ version 1.2.1. This estimation can be performed on the full, unpruned trees even when body mass data are missing for some tips. The ancestral state reconstruction yields body mass estimates at each internal node. For each branch, we then calculated the branch mass as the mean of the values at its two connecting nodes. We also discarded terminal branches for which tip body mass data were missing. Visualizations were implemented using the R packages ggplot2^107^ version 4.0.2 and ggdist^108^ version 3.3.3.

We employed similar methods to investigate whether morphological evolutionary rates correlate with body mass. Specifically, we performed PGLS on the terminal branch values, and used visualization to examine the overall pattern across all branches. For the randomly selected posterior trees, we summarized the unpartitioned, craniodental, axial, or appendicular rates using the RevGadgets^109^ version 1.2.1 and tidytree^110^ version 0.4.7 packages. The hexbin plots shown in Figures 4A-D were generated using the R package ggplot2^107^ version 4.0.2, hexbin^111^ version 1.28.5, and viridis^112^ version 0.6.5. For each of the 100 trees, we extracted the branches with rates in the upper 5% quantile as proxies for rate bursts, and compared their mass distribution against that of all branches.

