## Supplemental figures for "Constrained Body Mass Evolution and Decoupled Morphological Rates in Plesiosaurs"

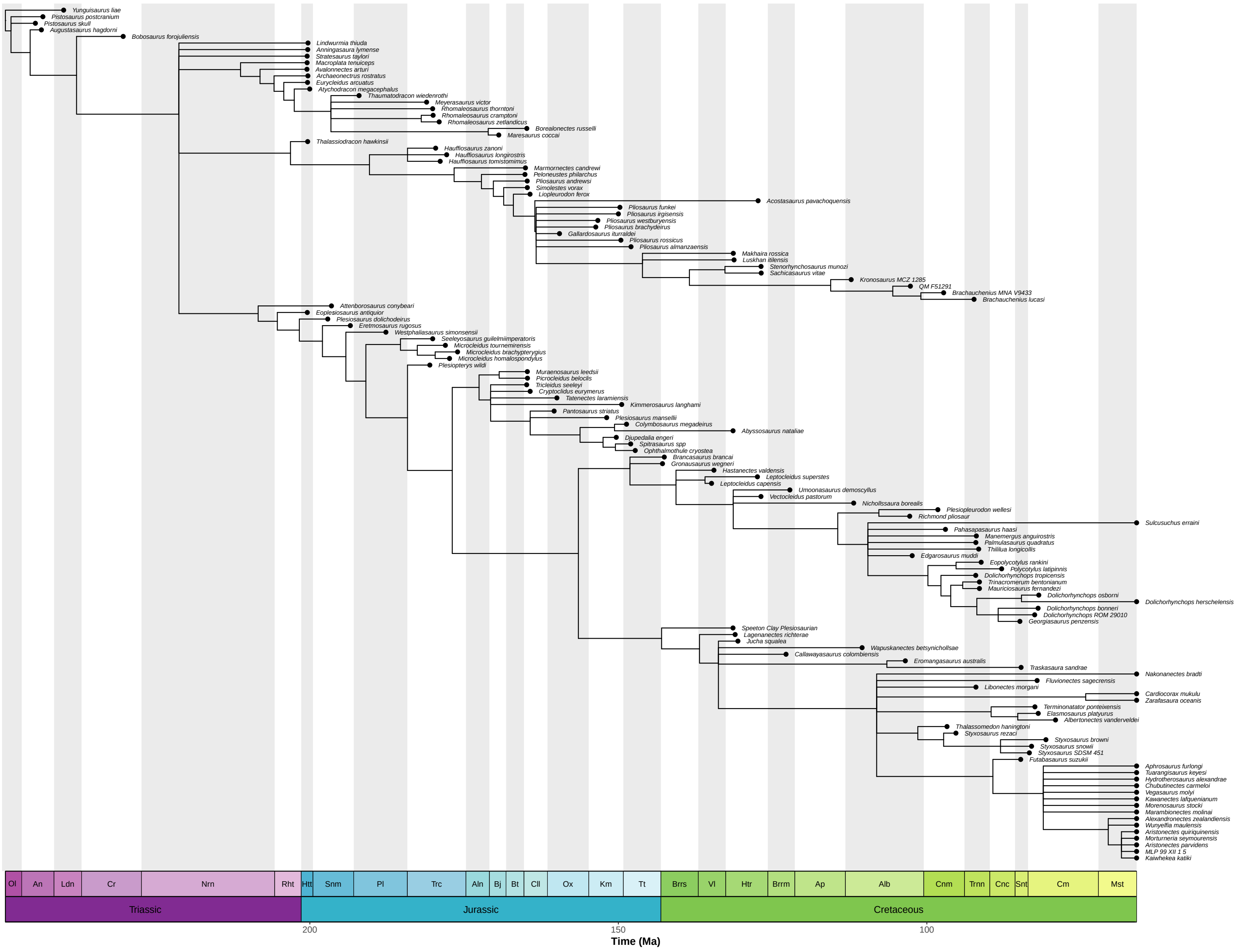


Figure S1: **50% majority-rule consensus tree of the skyline fossilized birth-death process.**

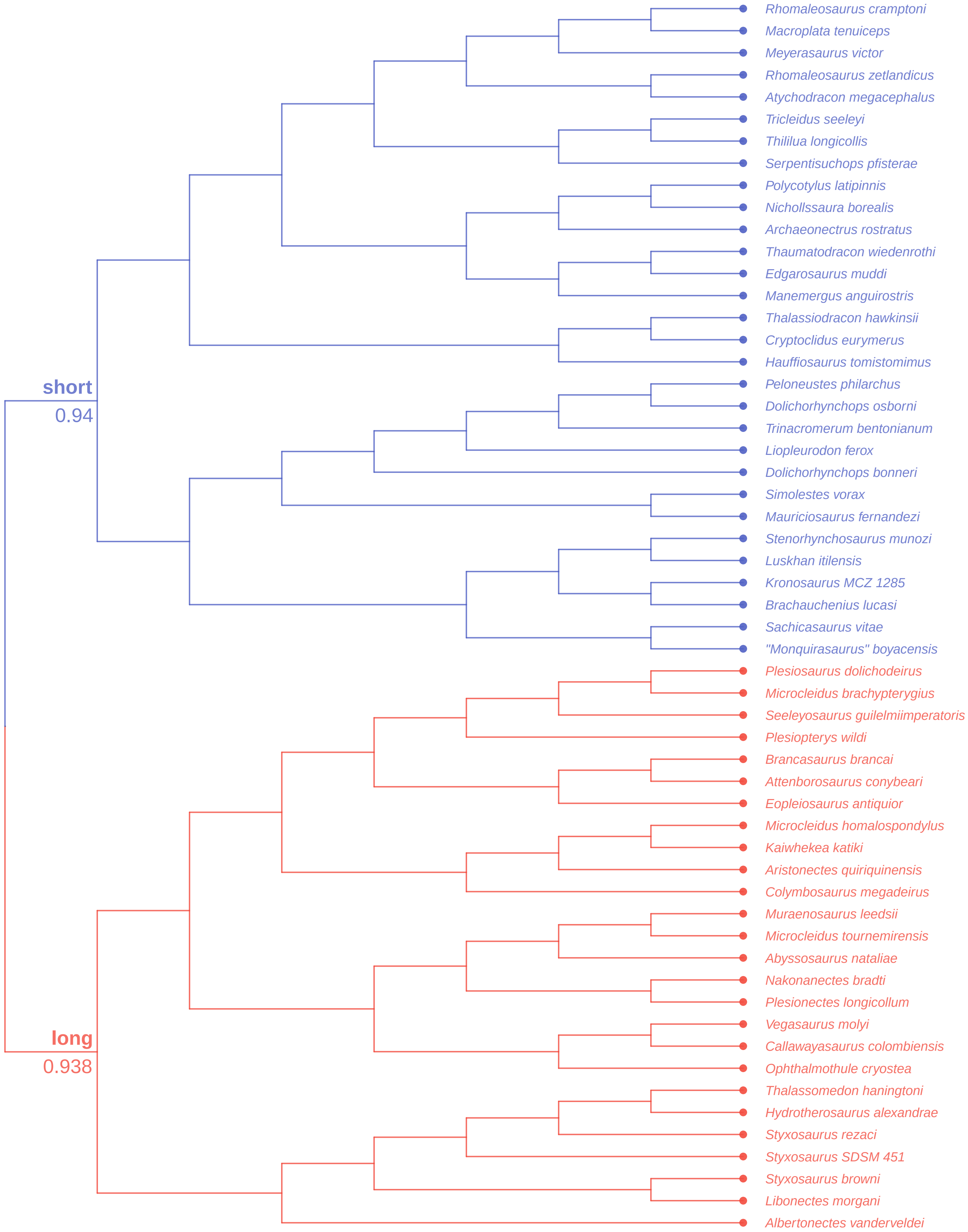


Figure S2: **Clustering dendrogram of plesiosaur neck length.** The values represent mean Jaccard similarities calculated based on bootstraping. Values larger than 0.85 indicate high stablility of clusters (see STAR Methods of the main text).
